# Lectin Microarray Analysis of Salivary Gland Glycoproteins from the Arboviral Vector *Aedes aegypti* and the Malaria Vector *Anopheles stephensi*

**DOI:** 10.1101/2024.07.10.602877

**Authors:** Ranjan Ramasamy, Xi Chen, Jian Zhang, Kokila Sivabalakrishnan, Sivasingham Arthiyan, Sinnathamby N. Surendran

## Abstract

Salivary gland glycoconjugates in human-feeding insects and arachnids can elicit hypersensitivity reactions. Glycoproteins introduced during blood feeding by arachnid ticks are responsible for red meat allergy or the α-gal syndrome. Dengue virus transmitted by *Aedes* mosquitoes incorporates mosquito glycoproteins into its envelope and malaria parasites transmitted by *Anopheles* mosquitoes reportedly express α-gal epitopes that are potentially derived from mosquito salivary glands. Tick salivary gland glycoproteins possess α-gal epitopes. We used microarrays containing forty different lectins to analyse salivary gland protein glycoconjugates from the principal arboviral vector *Aedes aegypti* and the rapidly spreading malaria vector *Anopheles stephensi*. Salivary gland glycoproteins of both mosquitoes possessed very similar lectin-binding specificities. Lectin-binding profiles showed the significant presence of oligomannose N-glycans and O-glycans, a limited presence of glycan structures capped with terminal GalNAc, GlcNAc, β linked Gal, α1-6 linked fucose, and no detectable sialic acids or terminal α-linked Gal in salivary gland glycoproteins.

## 1. Introduction

Mosquito and tick vectors transmit pathogens such as those respectively causing malaria and Lyme disease, and also elicit hypersensitivity reactions to salivary components inoculated into humans during blood feeding [1,2]. IgE antibodies to the trisaccharide epitope Galα1-3Galβ1-4GlcNAc-R (α-gal), initially elicited by α-gal epitopes in injected saliva, bind to α-gal from red meat and cause hypersensitivity termed the α-gal syndrome (AGS) or red meat allergy [2]. The clinical manifestations of AGS range from urticaria to fatal anaphylaxis [2]. Both arachnid ticks and insect mosquitoes are classified under the phylum *Arthropoda*. Mass spectrometry (MS) demonstrated the presence of α-gal in N-glycans of glycoproteins from the salivary glands (SGs) of two tick vector species, *Amblyomma americanum* and *Ixodes scapularis*, associated with eliciting AGS [3,4]. Enzymes with 1-3 α-galactosyl transferase (α1-3 GT) activity have been identified in *Ix. scapularis* [5]. The *Ix. scapularis* α1-3 GTs are homologous to mammalian α1-4 and β1-4 GTs but not mammalian α1-3 GT. Ixodes scapularis α1-3 GTs have close homologues in the principal arboviral mosquito vector *Aedes aegypti* and the major African malaria vector *Anopheles gambiae* [6] but α1-3 GT activities of the mosquito enzymes have not been investigated.

The presence of α-gal on the surfaces of mosquito SG-derived sporozoite stages of human and rodent malaria parasites (*Plasmodium falciparum* and *Plasmodium yoelii*/*berghei* respectively) was reported from the binding of Griffonia simplicifolia-I isolectin B4 (GSI-B4) and a monoclonal antibody against α-gal [7]. The absence of genes for galactosyl transferases in the Plasmodium genome [6,8], the absence of significant Golgi glycosylation [9], the failure to incorporate galactose from UDP-gal into glycolipids and glycoproteins in cultured asexual blood stages, and the non-specific binding of GSI-B4 to *P*. *falciparum* [8], raised the alternate possibility that *Plasmodium* sporozoites may have acquired α-gal containing glycoconjugates from the SGs of *Anopheles* mosquito vectors [6,10]. Dengue virus replicating in the arboviral mosquito vector *Aedes albopictus* derived C6/36 cell line reportedly acquired α-gal on virus envelope N-glycans based on lectin binding and MS analysis [11]. Detailed exoglycosidase and MS analysis of N- and O-glycans from whole larvae of *Ae. aegypti* and *An. gambiae*, however, showed no evidence of the presence of α-gal [12]. In the absence of prior characterisation of mosquito vector SG glycoconjugates, and conflicting reports on the presence of α-gal in mosquito tissues, we undertook microarray analysis with 40 lectins on SG proteins from *Ae. aegypti* and the rapidly spreading malaria vector *Anopheles stephensi* [13].

## 2. Methods

### 2.1 Preparation of mosquito salivary glands

Self-mating laboratory colonies of Ae. aegypti and An. stephensi were established and maintained as previously described [14,15]. Salivary glands from three to six-day-old non-blood-fed female mosquitoes were isolated using a dissecting microscope in 0.01M phosphate-buffered saline pH7.2 (PBS) at 0°C on glass microscope slides, and then stored at −80°C. One thousand salivary glands from *An. stephensi* and 700 salivary glands from *Ae. aegypti* collected in this manner were then freeze-dried and couriered to ZBiotech, USA for lectin microarray analysis. Maintenance and use of mosquito colonies for experiments was approved by the Animal Ethics Committee of the University of Jaffna (AERC/2023.03).

### 2.2 Protein extraction and lectin microarray analysis

Proteins from SGs were extracted with T-PER protein extraction reagent (Cat. No. 78510, Thermo-Scientific), and protein concentration determined using the Pierce BCA protein assay kit (Cat. No. 23225, Thermo-Scientific). Protein concentrations from the two SG preparations were equalized before biotin labelling with EZ-link NHS-LC-Biotin (Cat. No. 21336, Thermo-Scientific) followed by dialysis against PBS. Protein concentration after biotin labelling and moles of biotin per mole of protein were respectively 0.31 mg/ml and 5.02 mg/ml for *An. stephensi*, and 0.24 mg/ml and 4.62 mg/ml for *Ae. aegypti*.

The biotinylated cell lysate samples were then analysed with ZBiotech lectin arrays. The array features 40 plant or bacterial lectins immobilized on a microarray slide, as shown in Figure S1 and detailed in Table S1. Each slide was formatted into 16 subarrays. The carbohydrate-binding specificities of the lectins, assessed with glycan arrays, are shown in Table S2. The microarray slides were pretreated with lectin array assay buffer (137 mM NaCl, 2.7 mM KCl, 50 mM Tris, 0.1% Tween-20, supplemented with 2mM CaCl_2_ and MgCl_2_) at ambient temperature for 30 min. Biotin-labelled SG protein samples were then added at three different concentrations and incubated at ambient temperature for 1 h. After incubation, the microarray slides were washed and 0.2 μg/mL streptavidin Cy3 (Cat. No. 434315, Life Technologies) added. After incubation at ambient temperature for 1 h, the microarray slides were washed and scanned as summarized in Figure S2. Microarray slides were scanned at 532 nm at high-intensity settings (1 PMT). Scans revealed specific binding interactions in both SG protein samples, as illustrated in Figure S3. The scans were further analyzed with Mapix software (Innopsys), which determined binding signals by subtracting both the background signals and those from negative control probes, as shown in Figure S4.

## 3. Results

The nearly identical lectin binding specificities of *Ae. aegypti* and *An. stephensi* SG proteins by analysis of scans (Figure S4) showed that both mosquito vectors possessed similar SG protein glycoconjugate structures. Biotinylated SG proteins used at 5,10 and 20 µg/ml gave identical results, and the positive control amine-polyethylene glycol-Biotin and negative controls gave expected reactions (Figure S4). The ZBiotech lectin specificities (Table S2) were compared, where possible, with an independent recent determination [16] to reach a consensus on the main structural features in protein glycoconjugates present in SGs of both mosquito species, as summarised in Table 1.

**Table 1.** Structural features of salivary gland protein glycoconjugates from *Aedes aegypti* and *Anopheles stephensi*.

| Feature | Supportive lectin binding evidence |
| --- | --- |
| O-linked glycans containing core GalNAc | Strong binding to *Artocarpus integrifolia* (AIA) and weaker binding also to *Helix aspersa* agglutinin (HAA), *Helix pomatia* agglutinin (HPA), soyabean agglutinin (SBA), and *Vicia villosa* lectin (VVL) |
| N-linked oligomannose and/or hybrid glycans | Strong binding to *Calystegia sepium* lectin (Calsepa), concanavalin A (con A), *Galanthus nivalis* lectin (GNA), *Hippeastrum* hybrid lectin (HHL), *Lens culinaris* agglutinin (LCA), *Moringa oleifera* G lectin, *Narcissus pseudonarcissus* agglutinin (NPA) and mannose-specific recombinant prokaryotic lectin (RPL-Man2)<br>Strong binding to *Galanthus nivalis* agglutinin (GNA) and *Narcissus pseudonarcissus* agglutinin (NPA) suggests a high proportion of paucimannose N-glycan |
| Fucosylation | Strong binding to *Aleuria aurantia* lectin (AAL), *Lens culinaris* agglutinin (LCA), *Pisum sativum* agglutinin (PSA), *Ralstonia solanacearum* lectin (RSL), and absence of binding to *Psophocarpus tetragonolobus* agglutinin (PTA) and *Ulex europaeus* agglutinin 1 (UEA-1) suggests the presence of at least α1-6 linked fucose in the (GlcNAc)<sub>2</sub> core of N-linked glycans |
| Terminal βGalNAc, βGal, LacNAc | Strong binding to *Datura stramonium* agglutinin (DSA), *Lycopersicon esculentum* lectin (LEL), *Solanum tuberosum* lectin (STL), and weaker binding to *Dolichos biflorus* agglutinin (DBA), *Bauhinia purpurea* lectin (BPL), *Helix pomatia* agglutinin (HPA), soyabean agglutinin (SBA), *Vicia villosa* lectin (VVL), *Wisteria floribunda* agglutinin (WFA) |
| Terminal βGlcNAc | Strong binding to wheat germ agglutinin (WGA) |
| Terminal αGal not detected | No binding to *Griffonia simplicifolia*-I isolectin B4 (GSI-B4) and αGal-specific recombinant prokaryotic protein (RPL-αGal) |
| Terminal sialic acid not detected | No binding to sialic acid-specific recombinant prokaryotic protein (RPL-Sia2), *Sambucus nigra* agglutinin (SNA), and *Maackia amurensis* I and II lectins (MAL-I and MAL-II) |

## 4. Discussion

The findings from lectin microarray analysis of SG glycoproteins of *Ae. aegypti* and *An. stephensi* are broadly consistent with those obtained by a combination of exoglycosidase digestion and MS analysis of whole larval glycoproteins from *Ae. aegypti* and *An. gambiae* [12]. Common findings were the high oligomannose content, the presence of terminal βGalNAc, βGal, LacNAc and βGlcNAc, the absence of sialic acids, the absence of terminal αGal, and the detection of O-glycans [12], as well the presence of αFuc in N glycans of *An. gambiae* [17]. However, the terminal glucuronic acids and sulphate residues detected by MS [12] were not seen by lectin microarray analysis, and this may be due to the greater precision and sensitivity of MS detection, lack of lectins with requisite specificities and variation between SGs and other mosquito tissues. The α-gal detected on dengue virus envelope N-glycans when the virus is produced in *Ae. albopictus* C6/36 cells [11] may be due to the acquisition of α-gal containing glycoconjugates from foetal bovine serum used in the culture medium and requires further investigation. Analysis of N-glycans in saliva of the sandfly vector of visceral leishmaniasis, *Lutzomyia longipalpis*, by MS also showed a preponderance of oligomannose, fucosylation of the core GlcNAc, and terminal HexNAc residues, but O-glycans, terminal glucuronic acids and sulphate residues were not detected [18]. N-glycans in saliva from the vector of African trypmanosomiasis *Glossina morsitans morsitans* showed a preponderance of pauci-mannose and oligomannose, α-linked fucose in the core GlcNAc, and terminal GlcNAc in a study by MS [19]. O-glycans, terminal glucuronic acids and sulphate residues could not be detected in *Glossina morsitans morsitans* salivary proteins in this MS study [19]. Terminal αGal was not detected in protein glycoconjugates in the saliva of *L. longipalpis* [18] or *G. morsitans morsitans* [19]. This suggests that the presence of α-gal epitopes in the glycoproteins of arachnid tick vectors may not be a characteristic of insect mosquito vectors.

Oligomannose in N-glycans present in saliva of insect vectors when inoculated during blood feeding can bind a variety of C-type lectin receptors for mannose on human dendritic cells, macrophages and other immune cells as well as mannan binding protein in blood [20], and promote immune responses that can play a role in the varying types of hypersensitivity reactions to insect bites [1]. On the other hand, an absence of α-gal epitopes in the SGs in the two human insect vectors, in contrast to some tick vectors, suggests that insect bites may not play a role in red meat allergy or AGS. This requires to be established in more detailed studies. It is possible that α-gal was not detected in *Ae. aegypti* and *An. stephensi* SG proteins because it was present at levels below the threshold of detection in the lectin microarrays or because α-gal is possibly present only in mosquito glycolipids [21] which were not investigated in the present study. The previously reported detection of α-gal in *Plasmodium* sporozoites and SGs of anopheline vectors [7] may be caused by the non-specific GSI-B4 binding documented previously [8] and monoclonal antibody cross-reactivity, but requires further investigation because of its important implications for immunity against malaria [6-10].

## Data availability

All data used to draw conclusions are available in the article and its supplementary files.

## Abbreviations

α-gal: alpha-galactosyl epitope
α1-3 GT: 1-3 α-galactosyl transferase
AGS: alpha-gal syndrome
GSI-B4: *Griffonia simplicifolia*-I isolectin B4
MS: mass spectrometry
PBS: 0.01M phosphate-buffered saline pH7.2
SG: salivary gland

