## Supplementary Information for "Lectin Microarray Analysis of Salivary Gland Glycoproteins from the Arboviral Vector *Aedes aegypti* and the Malaria Vector *Anopheles stephensi*"

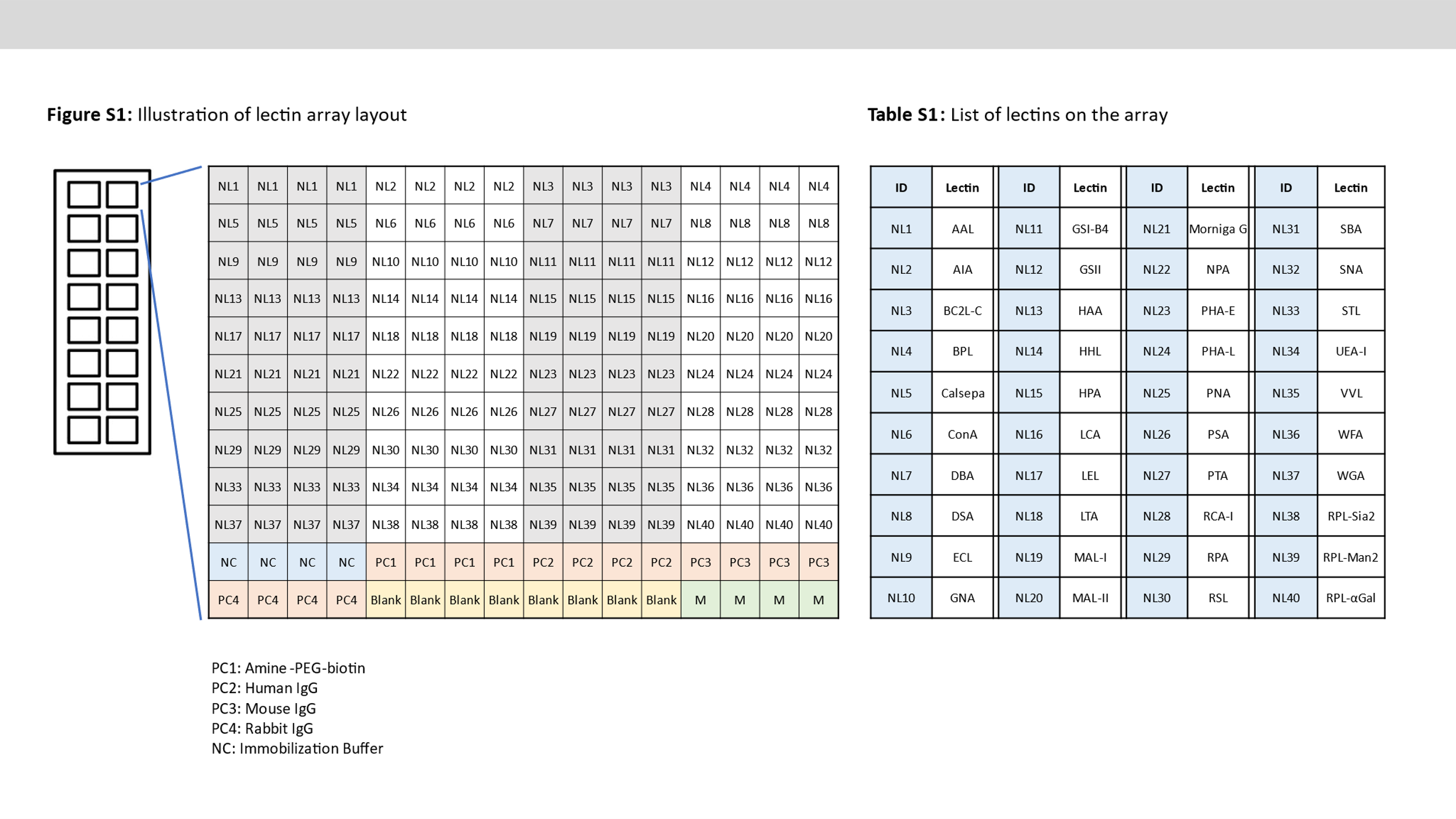

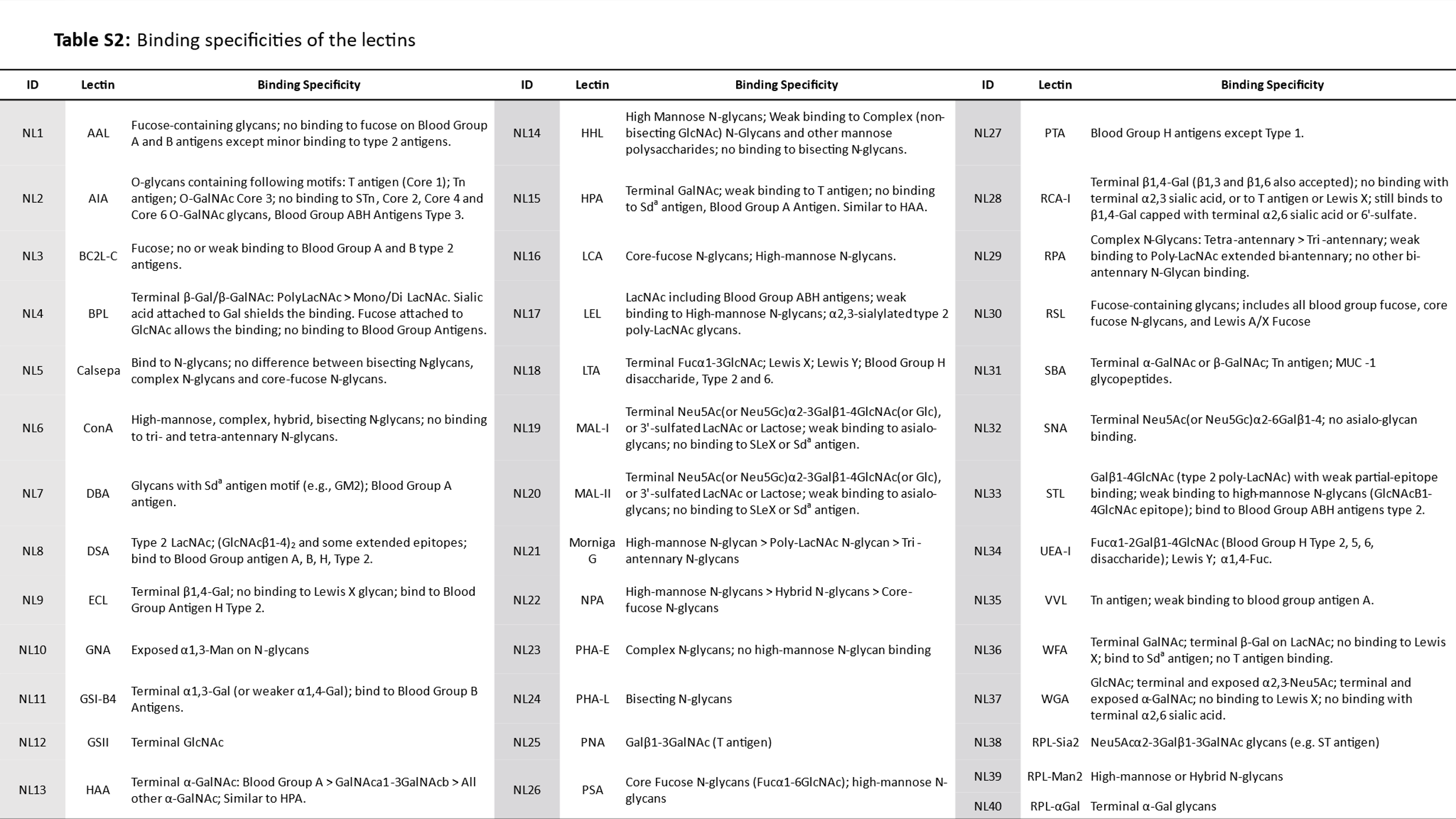

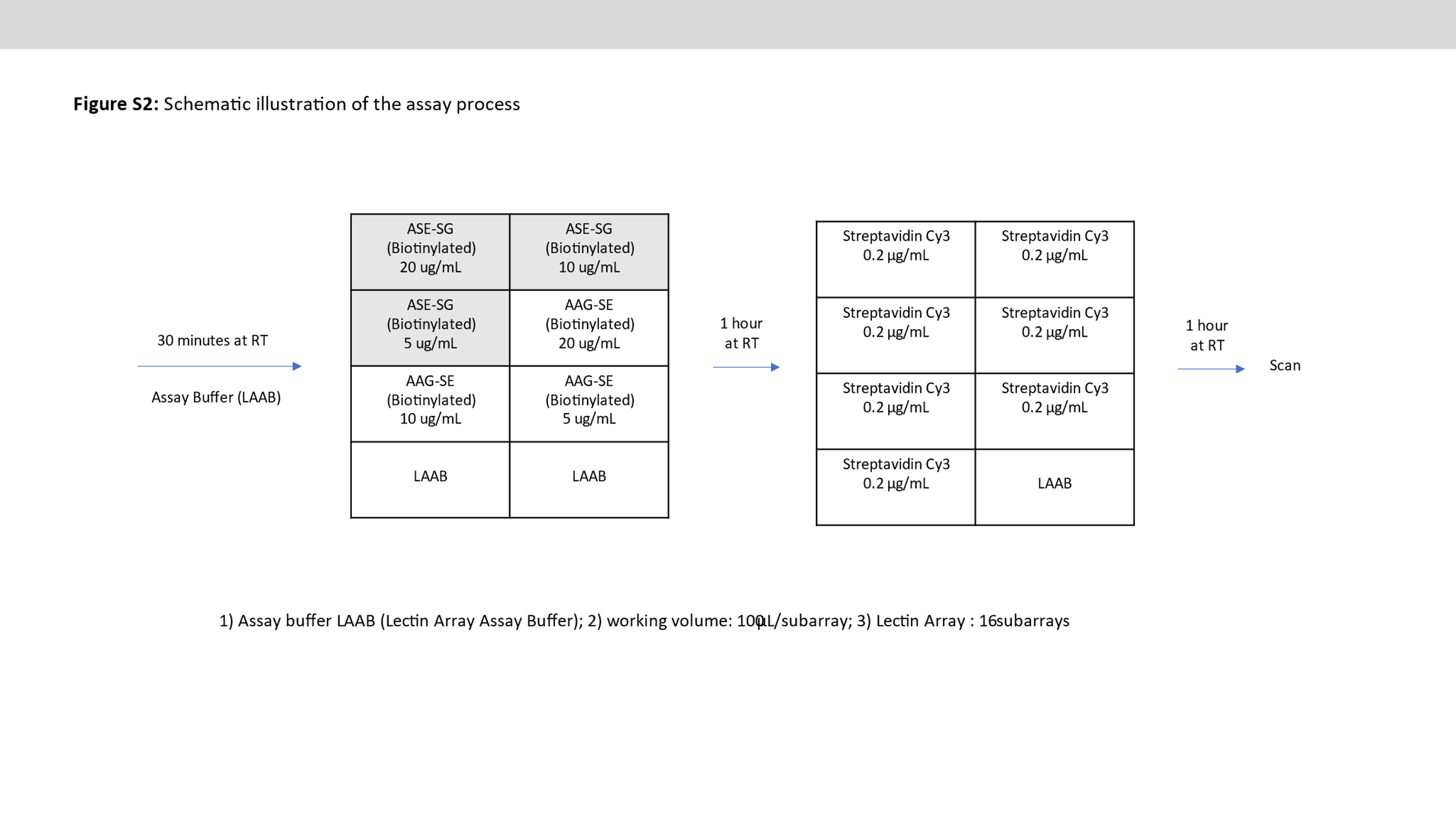

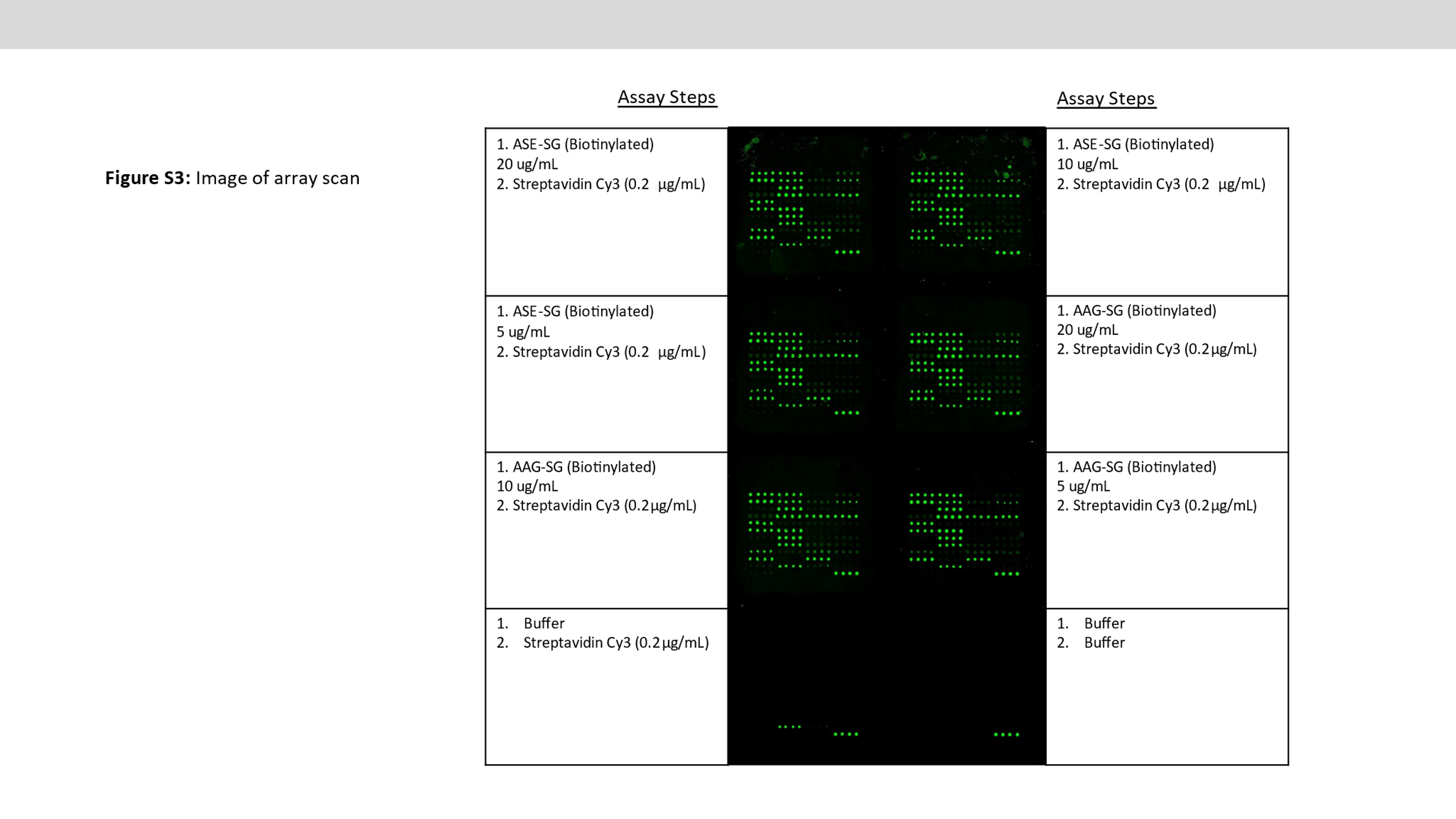

**Figure S4: Interactions of biotinylated SG extracts of *An. stephensi* and *Ae. aegypti* with various lectins**

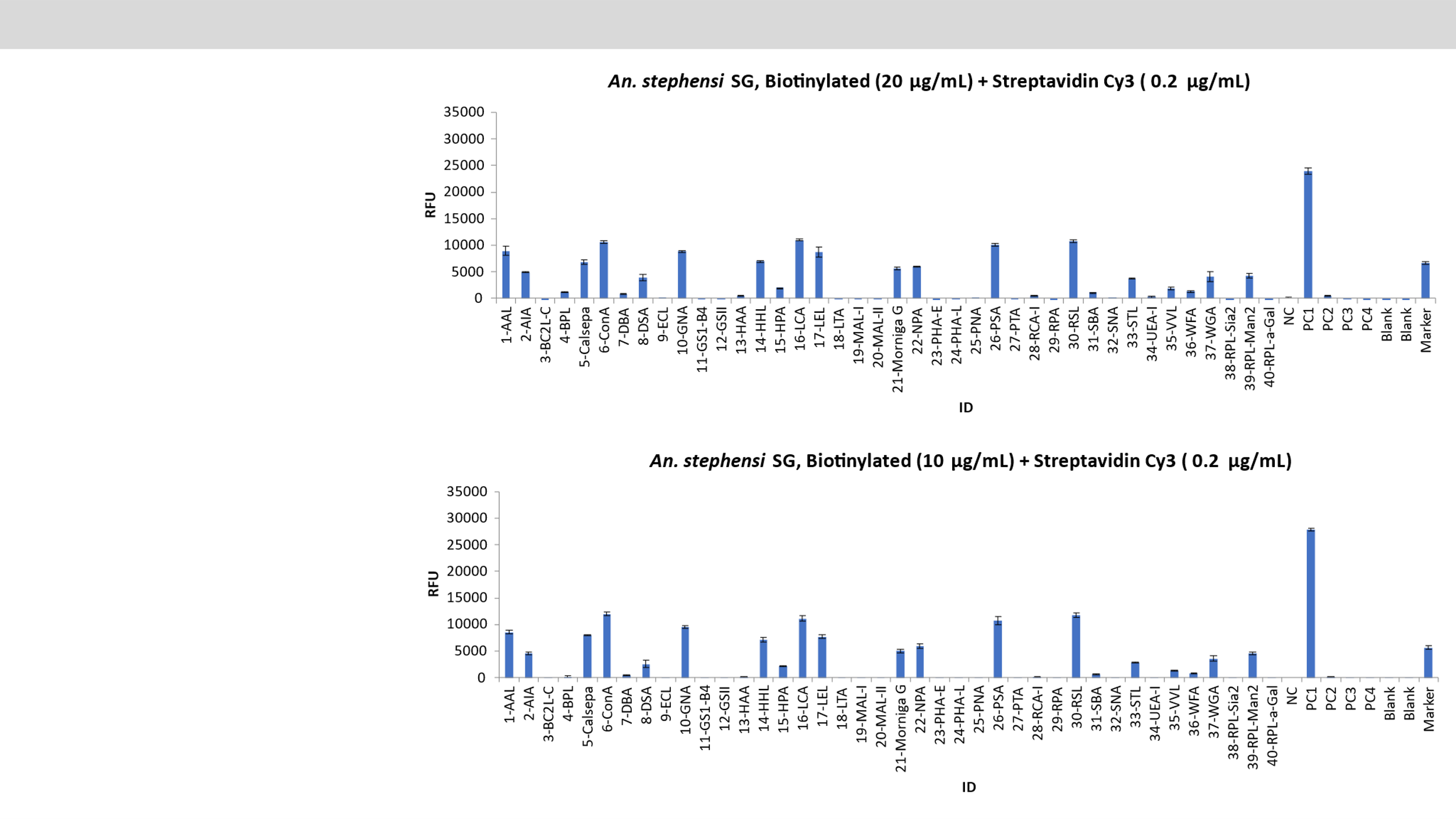

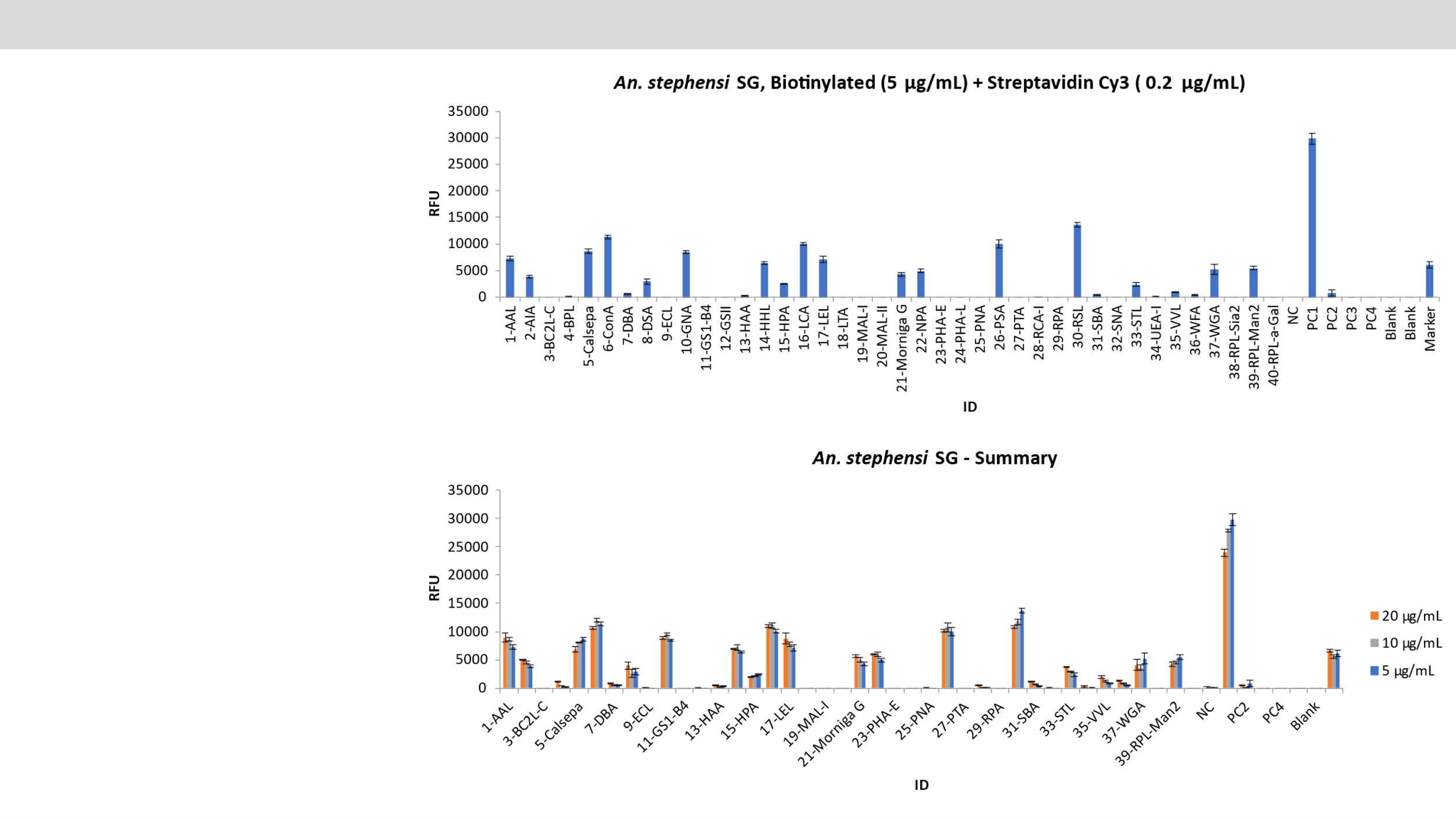

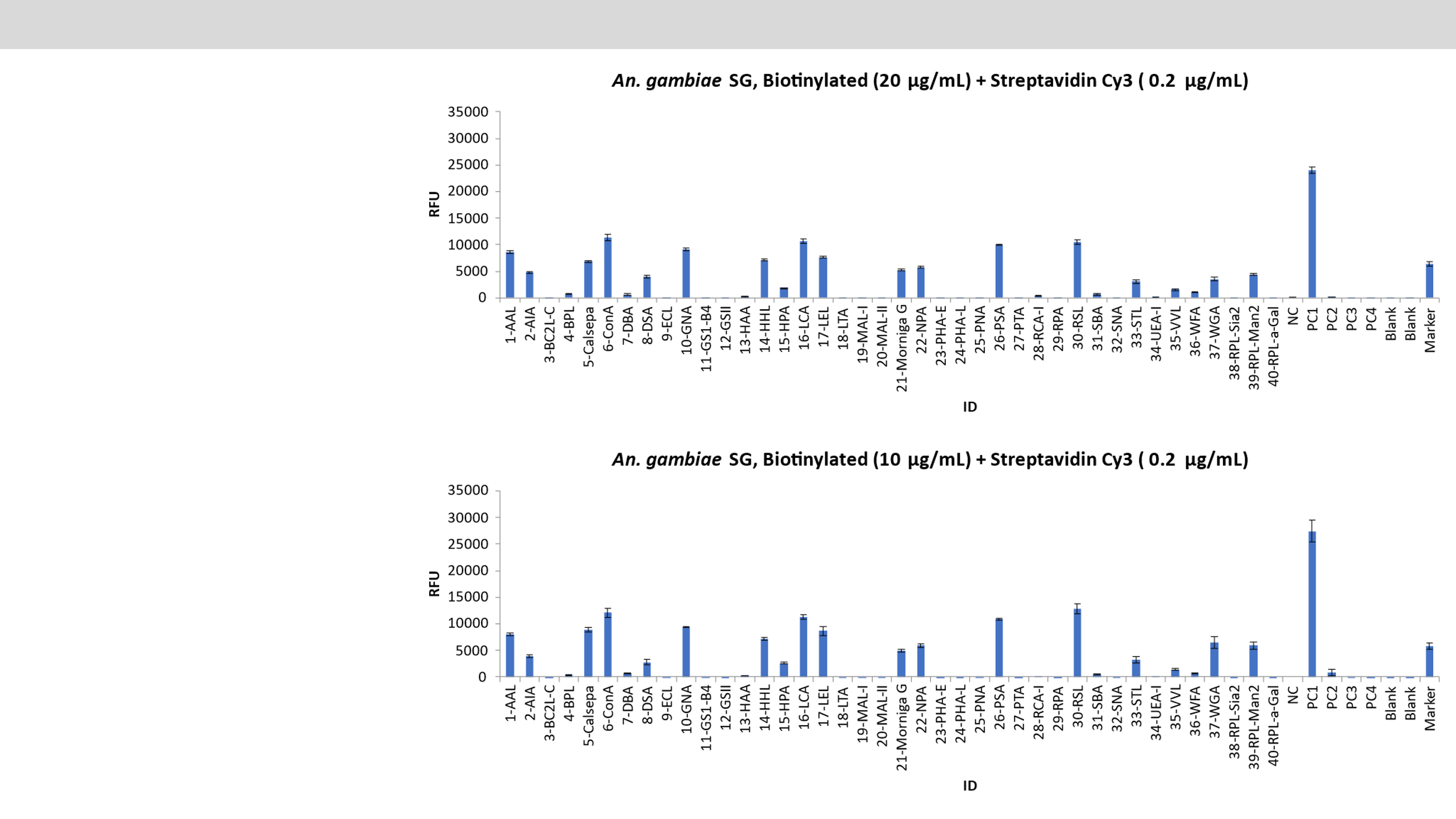

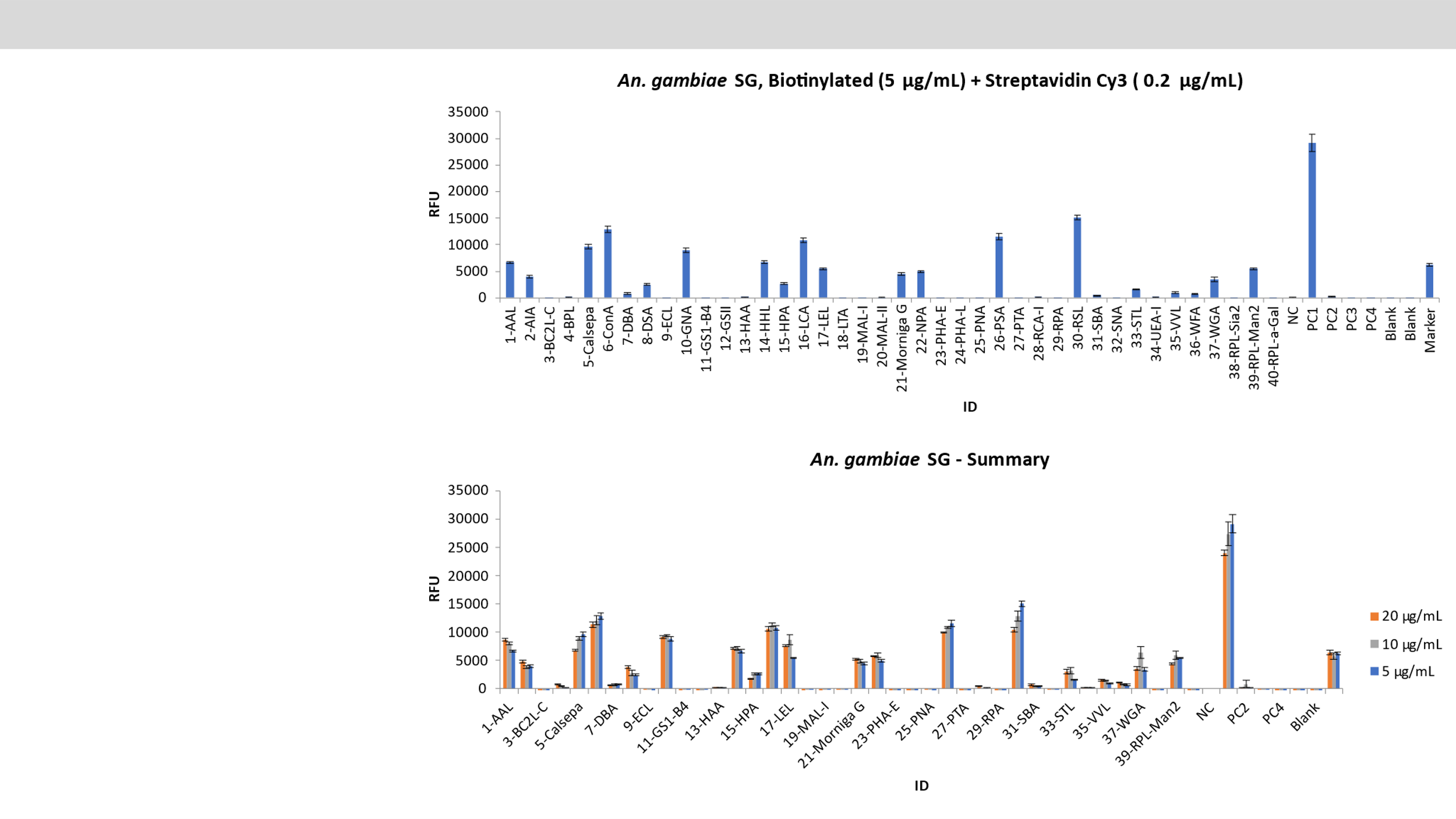

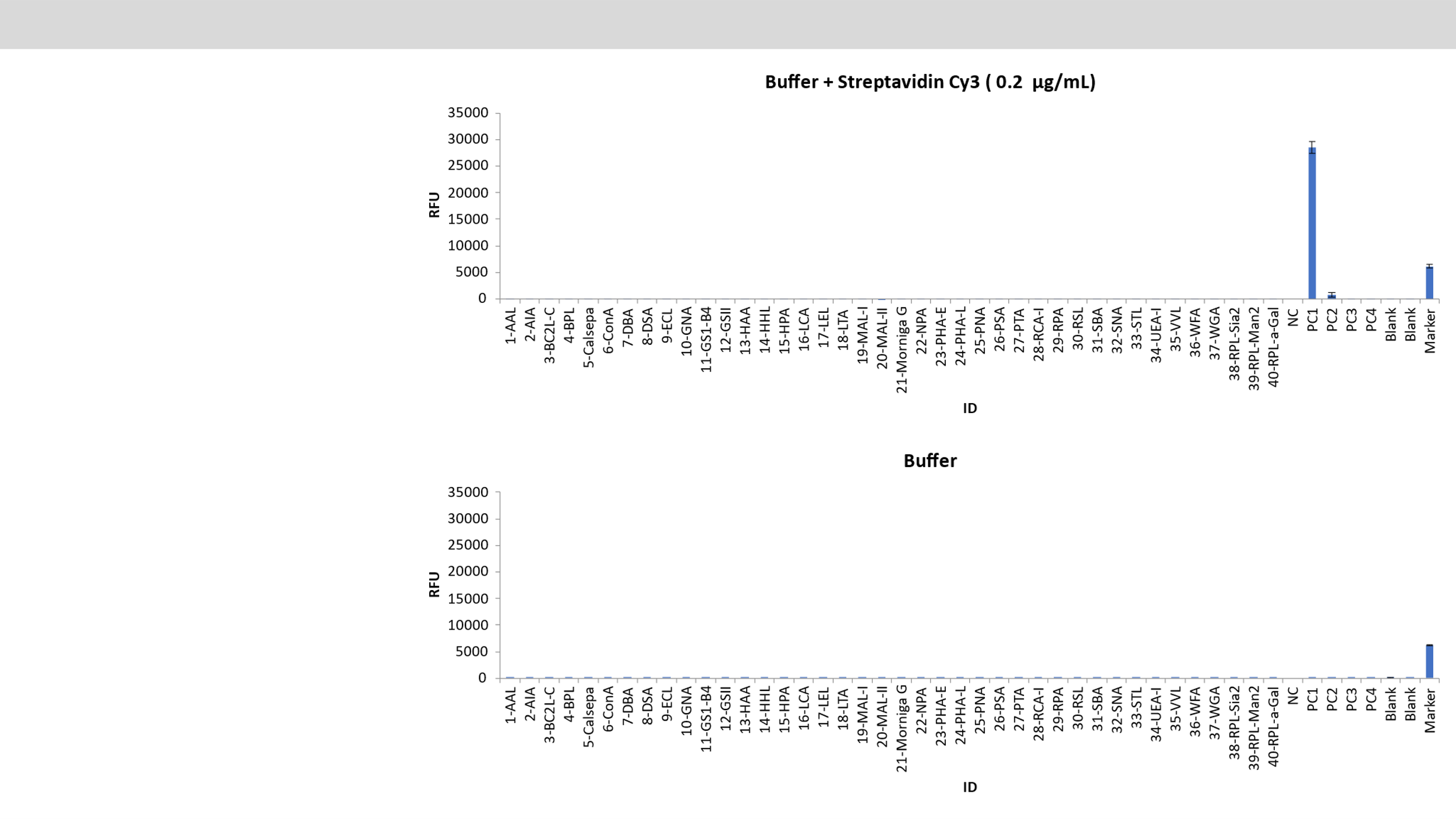
